# Functional genomics analysis identifies impairment of *HNF1B* function as a cause of Mayer-Rokitansky-Küster-Hauser syndrome

**DOI:** 10.1101/2022.04.26.489616

**Authors:** Ella Thomson, Minh Tran, Gorjana Robevska, Katie Ayers, Prarthna Gopalakrishnan Bhaskaran, Eric Haan, Silvia Cereghini, Alla Vash-Margita, Miranda Margetts, Alison Hensley, Quan Nguyen, Andrew Sinclair, Peter Koopman, Emanuele Pelosi

## Abstract

Mayer-Rokitansky-Küster-Hauser (MRKH) syndrome is a congenital condition characterized by aplasia or hypoplasia of the uterus and vagina in women with a typical 46,XX karyotype. This condition can occur as type I when isolated or as type II when associated with extragenital anomalies including kidney and skeletal abnormalities. The genetic basis of MRKH syndrome remains unexplained and several candidate genes have been proposed to play a role in its etiology, including *HNF1B*, *LHX1*, and *WNT4*. Here, we conducted a genomic analysis of 13 women affected by MRKH syndrome, resulting in the identification of candidate genes, including several novel candidates. We focused on *HNF1B* for further investigation due to its known association with, but unknown etiological role in, MRKH syndrome. We ablated *Hnf1b* specifically in the epithelium of the Müllerian ducts in mice, and found that this caused hypoplastic development of both the epithelial and stromal compartments of the uterus, as well as kidney anomalies, closely mirroring the MRKH type II phenotype. Using single-cell RNA sequencing of uterine tissue in the *Hnf1b*-ablated embryos, we analyzed the molecules and pathways downstream of *Hnf1b*, revealing a dysregulation of processes associated with cell proliferation, migration, and differentiation. Thus, we establish that loss of *Hnf1b* function leads to an MRKH phenotype, and generate the first mouse model of MRKH syndrome type II. Our results support the diagnostic value of *HNF1B* in clinical genetic testing for MRKH syndrome, and shed new light on the genetic causes of this poorly understood condition in women’s reproductive health.

## Introduction

MRKH syndrome (OMIM #277000) affects 1 in 4,500-5,000 women and features a spectrum of phenotypes, characterized by absent or incomplete development of the uterus, cervix and upper two-thirds of the vaginal canal in an otherwise typical 46,XX female. MRKH syndrome is classified into two main groups: type I, which is characterized by aplasia of uterus and/or vagina, and type II, which is associated with additional malformations. A subset of type II consists of Müllerian duct aplasia, renal aplasia, and cervicothoracic somite dysplasia (MURCS) association (OMIM #601076) (Duncan 1979). The most common extragenital malformations include renal (30-57% of cases), skeletal (30-44%), auditory (10-25%), and less frequently, heart anomalies (Morcel 2007; Rall 2015a). However, ovarian development and function are usually unaffected, allowing conception of biological children through gestational surrogacy or uterine transplantation (Beski 2000, Brännström 2015, Brännström 2016). Because women with MRKH syndrome develop otherwise typical female secondary sex characteristics, most are not diagnosed until adolescence due to primary amenorrhea. The diagnosis typically causes significant psychological distress for these women and their families (Carroll 2020).

The genetic cause(s) of MRKH syndrome remains unknown. The situation is complicated by the variability of phenotypes, and the polygenic and multifactorial nature of the condition (Wottgen 2008, Duru 2009, Rall 2015b, Milsom SR 2015). As a result, it has not been possible to develop diagnostics, or satisfactory genetic counselling strategies for affected women regarding the cause or nature of their specific condition or the potential risk for their future children.

Both sporadic and familial cases have been reported, and analysis of familial clustering have suggested the existence of genetic predisposition (Herlin 2016, Herlin 2014, Ma 2016). Also, both autosomal recessive, and autosomal dominant patterns – often characterized by incomplete penetrance and variable expressivity – are known to occur (Shokeir 1978, Herlin 2014; Wottgen 2008, Demir Eksi 2018). To date, more than 50 genes have been proposed as candidates for MRKH syndrome, including *RBM8A*, *WT1*, *TBX6*, and *SHOX* (Jacquinet 2016, Mullen 2014, Gervasini 2010, Fontana 2017). However, due to the lack of functional studies, the specific role of these proposed candidate genes in Müllerian duct (MD) development is still unclear. Consequently, most gene changes associated with MRKH syndrome are classified as variants of uncertain significance and their diagnostic potential remains undefined.

In addition to clinical investigations, animal studies have identified a number of genes that are involved in MD development and that can also be considered potential candidates for a role in MRKH syndrome. *Wnt4* was found to be necessary for urogenital development in the mouse (Vainio 1999), and mutations of *WNT4* in humans have been associated with uterine malformations (Biason-Lauber 2004). However, due to hyperandrogenism and virilization of the phenotype, this has been proposed as a separate entity from MRKH syndrome (OMIM #158330) (Biason-Lauber 2007). In addition, other genes known to play critical roles in MD development of the mouse have yet to be associated with MRKH syndrome. These include *Pax2*, *Digh1*, Dach1/2, *Hoxa11*, and *Wnt5a* (Jacquinet 2016, Mullen 2014).

Several human chromosomal imbalances have been described in MRKH syndrome, including 17q12, 16p11.2, and 22q11.21 (Sundaram 2007, Cheroki 2008, Nik-Zainal 2011, Ledig 2018). Deletions of 1.4-1.8 Mb in 17q12 represent the most frequent chromosomal rearrangements, accounting for up to 9% of MRKH cases (Edghill 2006, Bernardini 2009). This region contains the genes *LHX1* and *HNF1B*, which are both expressed in the epithelium of the developing genitourinary tract (Kobayashi 2004, Coffinier 1999). Ablation of *Lhx1* in the mouse results in the block of MD elongation and uterine hypoplasia, demonstrating the critical role of this factor in the development of the female reproductive tract (Kobayashi 2004, Huang 2014). *Hnf1b* has been shown to be an important factor for kidney function and development (Heliot 2013, Massa 2013, Lokmane 2010), and *HNF1B* mutations have been associated with genitourinary malformations including MRKH women (Fontana 2017, Lindner 1999, Oram 2010). However, no specific role for *Hnf1b* in reproductive tract development has been established experimentally.

In the present study, we performed SNP microarray and whole exome sequencing in a cohort of 13 women with MRKH syndrome and 11 unaffected family members, and identified a number of candidate genes. Among them, *HNF1B* was selected for *in vivo* functional analysis due to the known association of 17q12 deletions with MRKH syndrome and the need to establish a functional contribution of *Hnf1b* to the pathophysiology of MRKH. By ablating *Hnf1b* specifically in the MD epithelium in mice, we generated a novel model of MRKH type II, characterized by uterine hypoplasia associated with kidney anomalies. Using single-cell RNA sequencing, we identified the molecules and pathways that were affected by *Hnf1b* ablation in the main cell populations of the uterus. Our translational approach shows that loss of *HNF1B* disrupts MD development and uterine differentiation, revealing its role in MRKH syndrome.

## Materials and Methods

### Participants

Inclusion criteria for recruitment of subjects to this study were MRKH type I or II diagnosed clinically or radiographically. For two participants, we were unable to retrieve information related to their specific type of MRKH syndrome. Participants affected by MRKH syndrome, and (where possible) their unaffected family members, were recruited by their primary clinician. Each participant was de-identified and assigned a case number (e.g., MRKH01, MRKH02, etc.). Written informed consent was received prior to participation.

### Genetic Analysis

DNA was extracted from peripheral blood by the Victorian Clinical Genetics Services (VCGS) Molecular Genetics laboratory or local laboratories using standard protocols. DNA quality was assessed by a TapeStation 2200 (Agilent) and concentration was measured using Qubit dsDNA Broad Range (Thermofisher). Cytogenetics confirmed that all women had a 46,XX karyotype. Extracted DNA samples were sent for SNP microarray and whole exosome sequencing. When available, DNA of unaffected family members was also analyzed using whole exome sequencing.

### SNP microarray

Samples were analysed for copy number variants (CNVs) by SNP microarray undertaken by the VCGS cytogenetics laboratory using the Omni 2.5-8 (Illumina) technology. This platform provides high-level SNP coverage of exonic regions (1–2 SNP probes per 10 kb) through the entire genome. The assay detects copy number changes to an average resolution of 10–20 Kb, long continuous stretches of homozygosity (LCSH) regions >2 Mb, and mosaicism between 10– 20%. Library preparation and sequencing was carried out according to the manufacturer’s instructions.

### Whole exome sequencing

Whole exome sequencing was performed by the VCGS Translational Genomics Unit using the Agilent SureSelect QXT (transposase-based fragmentation) Clinical Research Exome (CRE) technology with 100× coverage on an Illumina HiSeq2500 instrument. Library preparation, sequencing, and analysis were carried out as per the manufacturer’s instructions. Variants were characterised applying the Cpipe (ver. 0.9.8) bioinformatic detection pipeline (http://cpipeline.org) (Sadedin et al., 2015). An internal database was used to filter technical and locally recurrent variants against participant data. Variant population frequency data was accessed from both GnomAD and ExAC. The predicted effect on gene function was calculated by the in-silico software programs SIFT (http://sift.bii.a-star.edu.sg/), Polyphen2 (http://genetics.bwh.harvard.edu/pph2/), CADD raw scores (Kircher et al., 2014), and MutationTaster2 (Schwarz 2014). Gene-associated phenotypes were also assessed by searching human cases reported in OMIM (http://www.omim.org/), mouse models in the Mouse Genome Informatics database (http://www.informatics.jax.org/) and known roles in physiological processes.

### Mouse strains and husbandry

*Hnf1b^fl/fl^* mice were donated by Dr S. Cereghini (Heliot 2013), and *Wnt7a-Cre* mice were a generous gift of Dr R. Behringer (Huang 2014). ROSA26-LacZ (B6;129S-Gt(ROSA)26Sor/J) mice were obtained by the Jackson Laboratory (Friedrich 1991). All mouse strains were maintained on a C57BL/6 genetic background. Samples were collected at 12.5- and 13.5-days post coitum (dpc), 0- and 7-days post-partum, and 4 months (4M) of age. All animal procedures were approved by the University of Queensland Animal Ethics Committee.

### RNA expression analysis

Samples were stored in RNAlater (Invitrogen) and total RNA was extracted using either an RNAeasy mini or RNAeasy micro kit (Qiagen). Following RNA amplification into cDNA using a high-capacity cDNA reverse transcription kit (Applied Biosystems), real-time PCR assays were performed on three biological replicates per genotype using a SYBR Green PCR master mix (Applied Biosystems). Quantitative real-time PCR was performed using a QuantStudio 3 system (Applied Biosystems). Expression was normalized to Tbp. Primers used are listed in Suppl Table 1.

**Table 1.** Summary of variants identified by SNP microarray.

| **Patient** | **MRKH Type** | **CNV** | **Size** | **Coordinates** |
| --- | --- | --- | --- | --- |
| MRKH01 | II | Duplication: 1q21.1<br>Deletion: 17q12 | 0.37 Mb<br>1.4 Mb | 145,394,955 – 146,762,959<br>34,815,551 – 36,220,373 |
| MRKH03 | II | Duplication: 3q13.12q13.13 | 0.4 Mb | 107,822,130 – 108,214,218 |
| MRKH05 | I | Duplication: 17q25 | 0.7 Mb | 79,857,201 – 80,583,230 |
| MRKH06 | I | Duplication: 2q24.2 | 0.5 Mb | 160,822,423 – 161,294,055 |
| MRKH08 | II | LCSH: Chr4 | >5 Mb | 140,586,701 – 145,900,002 |
| MRKH10 | II | Deletion: Xp22.33<br>LCSH: Chr11 | 0.15 Mb<br>>5Mb | 167,755 – 182,276<br>101,153,216 – 107,428,694 |

LCSH = long continuous stretches of homozygosity.

### Histological preparation and immunohistochemistry

Samples were fixed overnight in 4% paraformaldehyde-phosphate buffered saline (PBS). After fixation samples were embedded in paraffin and sectioned at 5 μm. For immunohistochemistry, ovary sections were deparaffinized in xylene and rehydrated using sequential washes and decreasing alcohol concentrations. Heath-mediated antigen retrieval was performed in 10 mM citrate buffer or Tris-based buffer (Vector). Immunofluorescence was done as previously described (Pelosi 2013) using the following primary antibodies: CDH1 (1:100, BD Biosciences #610182), HNF1B (1:100, Thermo Fisher Scientific #PA-50531), FOXL2 (1:100, Novus Biologicals #NB100-1277), PAX2 (1:100, BioLegend #901001), PH3 (1:100, Abcam #5176), LAMA (1:100, Sigma #L9393), VIM (1:200, Abcam #92547), SMA (1:100, Sigma #A2547).

Images were taken on a LSM710 confocal microscope (Zeiss) and processed using Adobe Photoshop.

### Single-cell RNA sequencing

Dissected reproductive tracts were cut into small pieces and incubated in a solution of DMEM/F12/Collagenase/Hyaluronidase/FBS (Stemcell) for 60 min at 37C. Following centrifugation at 300 x g for 5 minutes, cells were resuspended in 0.25% Trypsin-EDTA and incubated for 5 min. Hanks’ Balanced Salt Solution Modified (HBSS, Stemcell) supplemented with 10% FBS was added. After centrifugation, cells were resuspended in 1 mg/mL DNase I (Stemcell) under continued but gentle pipetting for 1 minute. Cells were washed three times in HBSS, resuspended in PBS, and cell count was performed to determine viability and cell concentration. Single cell suspension was partitioned and barcoded using the 10X Genomics Chromium Controller (10X Genomics) and the Single Cell 3’ Library and Gel Bead Kit v3.1 (V3.1; 10X Genomics; PN-1000121). The cells were loaded onto the Chromium Single Cell Chip G (10X Genomics; PN-1000120) to target 10,000 cells. GEM generation and barcoding, cDNA amplification, and library construction was performed according to the 10X Genomics Chromium User Guide. The resulting single cell transcriptome libraries contained unique sample indices for each sample. The libraries were quantified on the Agilent BioAnalyzer 2100 using the High Sensitivity DNA Kit (Agilent, 5067-4626). Libraries were pooled in equimolar ratios, and the pool was quantified by qPCR using the KAPA Library Quantification Kit - Illumina/Universal (KAPA Biosystems, KK4824) in combination with the Life Technologies Viia 7 real time PCR instrument. Denatured libraries were loaded onto an Illumina NextSeq- 500 and sequenced over 2 runs using a 150-cycle High-Output Kit as follows: 28bp (Read1), 8bp (i7 index), 111bp (Read2). For additional sequencing, denatured libraries were loaded onto an Illumina NovaSeq6000 and sequenced using one SP 100 cycle flow cell and one S1 100 cycle flow cell as follows: 28bp (Read1), 8bp (i7 index), 91bp (Read2). Read1 supplies the cell barcode and UMI, i7 the sample index, and Read2 the 3’ sequence of the transcript. Data pre- processing was performed using cellranger 3.0.2 (10x Genomics) and a mouse GRCm38 reference. Final read depth was ∼18,000 – 26,000 reads per cell across all samples. Library preparation and NextSeq sequencing were performed at the Institute for Molecular Bioscience Sequencing Facility (University of Queensland). NovaSeq sequencing was performed at Microba, based at the Translational Research Institute in Brisbane.

### Analysis of single-cell RNA sequencing

Raw sequencing data in BCL format were processed into the single-cell feature counts using Cell Ranger software, version, from 10X Genomics. For the downstream analysis, we used Seurat package version 4.0.3 to load the six Cell Ranger count matrices into Seurat objects. Cells with fewer than 200 genes and more than 5% of the reads mapped to mitochondrial were filtered during the preprocessing step. To remove batch effects, each dataset was normalized with SCTranform (Hafemeister 2019). We used the function SelectIntegrationFeatures() to choose the top 5000 variable genes across six datasets prior to running the FindIntegrationAnchors() function to identify integration anchors. These anchors were subsequently used to merge all six samples into one Seurat object using IntegrateData() function.

The integrated data was then used for dimensionality reduction and cell clustering, which used the top 5000 most variable genes of the integrated data. Dimensionality reduction was performed using Principal Component Analysis (PCA) to project the high dimensional space with 5000 genes to the reduced space of 50 PCs. Uniform manifold approximation and projection (UMAP) plot was generated using the first 30 PCs as input. 30 PCs were determined as a good threshold by using the ElbowPlot function (Suppl Figure 1). The visualization of the integrated data with cells was produced (Suppl Figure 2). Next, we clustered the cells using the Louvain algorithm (Traag (2019) by running the FindNeighbors() and FindClusters() functions. Using the resolution 0.02 for the FindClusters() function, we identified 9 clusters of cell types from the integrated data. Differential expression analysis was performed using the Wilcoxon Rank Sum test to find the top differentially expressed genes of each cluster. Clusters annotation was based on differentially expressed genes as described in the results section. Stromal cells subclusters were determined by subsetting the respective clusters, rescaling the data and repeating unsupervised clustering on the subset of the larger Stromal Cells cluster. We obtained four subclusters using a higher resolution parameter when running the FindClusters() function. Markers for the clusters were found by performing differential expression analysis using Wilcoxon Rank Sum and the cluster annotation was then done based on the gene markers. Differential gene expression between conditions was performed with an “pseudo-bulk” expression approach (Tung 2017). For each cell type, the pseudo-bulk expression profile was generated by summing the gene counts of all cells with the same condition (controls and *Hnf1b* mutants). Genes with an expression lower than the threshold of log count per million were filtered. To normalize library size differences for each pseudo-bulk samples, we used the calcNormFactors() function from EdgeR (Robinson 2010). The normalised pseudo-bulk expression of each sample was weighted to the condition from which it originated and fed to a linear model for differential expression analysis. Finally, empirical bayes statistic through eBayes() function was applied to rank the genes in order of significance (Law 2014).

**Figure 01.**
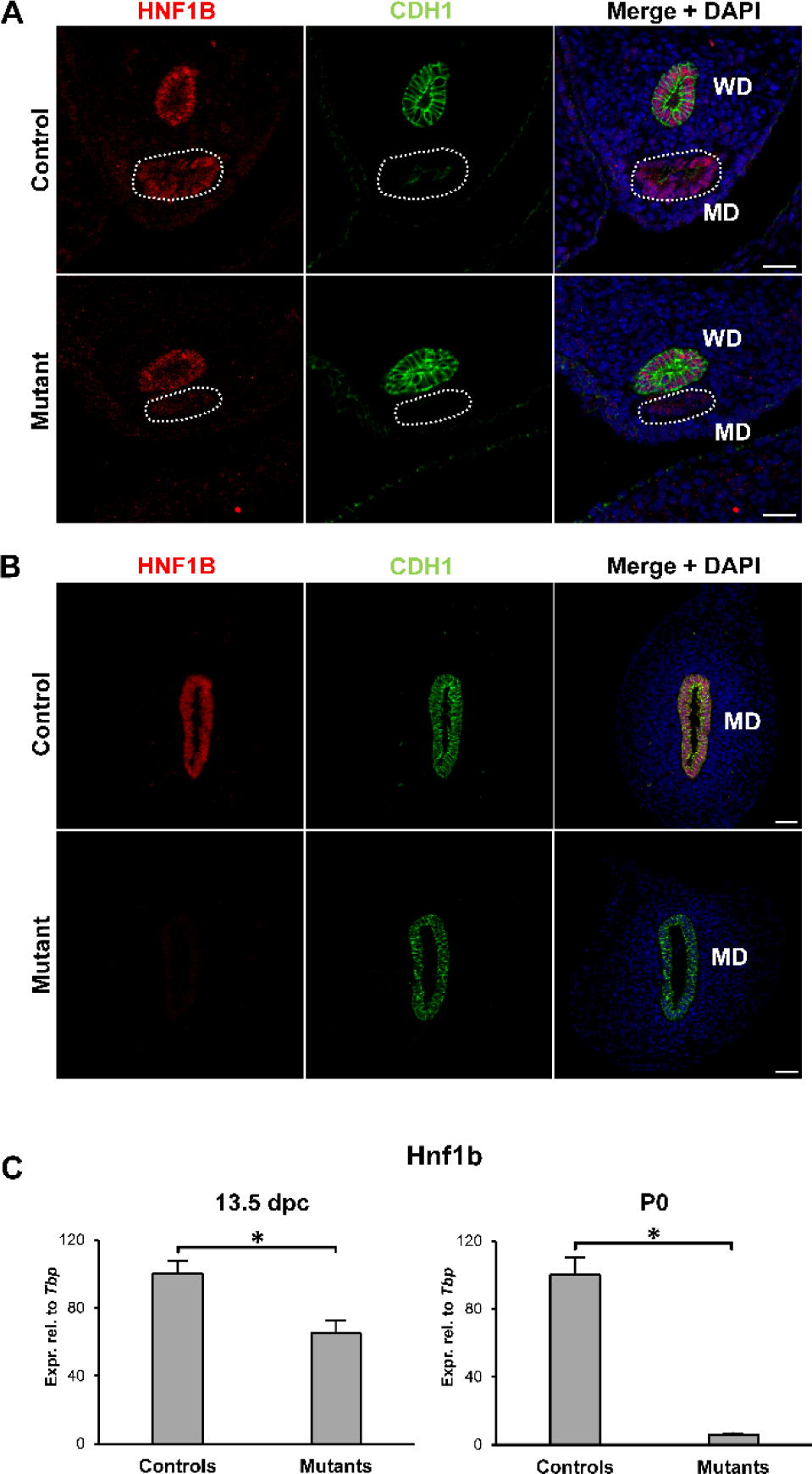
Hnf1b ablation in Mullerian duct epithelium. (A) In control animals, HNF1B was expressed in both the Wolffian ducts (WD) and the Mullerian ducts (MD, dashed line) at 13.5 dpc. In mutants, HNF1B expression was lost specifically in the MD. (B) At P0, only the MD remains in female animals, and immunohistochemistry showed no expression in the mutants. The epithelial marker CDH1/E- cadherin was used to visualize the WD at 13.5dpc and the MD at P0. Scale bar = 25 μm. (C) Real-time PCR of whole mesonephric tissues showed a reduction in *Hnf1b* expression at 13.5 dpc in mutant samples, consistent with the presence of the *Hnf1b*-expressing WD. At P0, expression of *Hnf1b* was lost in the mutants. * = p < 0.05.

**Figure 02.**
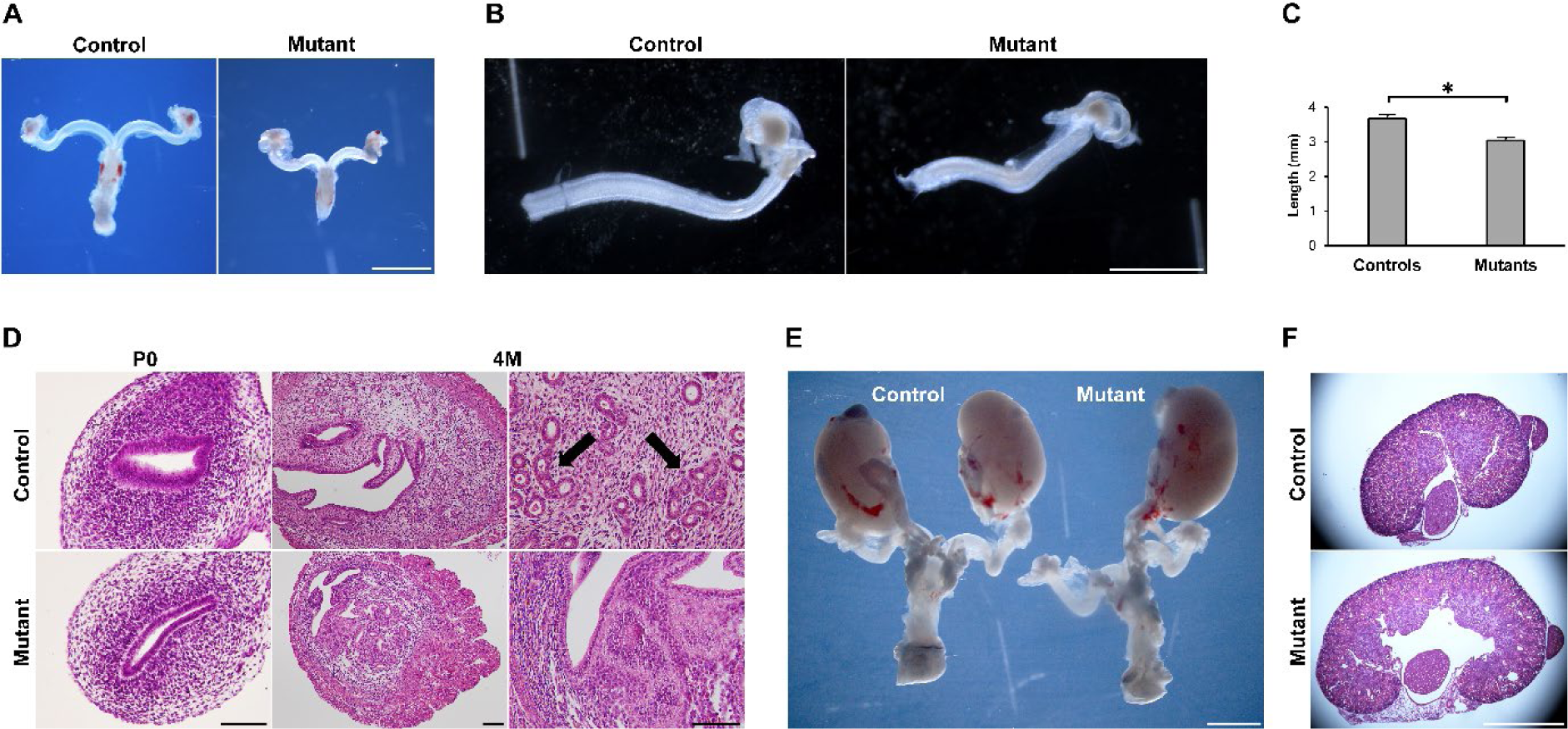
Phenotype of the Hnf1b mutant genitourinary system. (A-B) Gross morphology of reproductive tracts showed uterine horns of Hnf1b mutant mice that appeared smaller than controls. (C) Measurement of the uterine horn length at P0. (D) Hematoxylin and Eosin staining at P0 and 4M. Both epithelial and stromal compartments of the endometrium appeared hypoplastic in the mutant uteri. 4M control samples showed uterine glands (arrows) that were lacking in the mutants. (E) Unilateral kidney agenesis in a mutant animal. (F) Hematoxylin and Eosin staining of kidney samples showing a case of cystic kidney in the mutant. Scale bars = 2 mm (A, E, F), 1 mm (B), 20 μm (D, P0 and 4M right), 200 μm (D, 4M left). * = p < 0.05.

### Statistical analyses

For gene expression using real-time PCR, statistical analysis was performed by the unpaired t-test (GraphPad). Data were represented as mean expression levels with standard error of the mean (SEM). For comparison of unilateral kidney agenesis occurrence, significance was calculated using the Fisher’s exact test.

## Results

### Identification of copy number variants by SNP microarray

In our cohort, SNP microarray analysis identified copy number variants in six participants, two affected by MRKH type I (MRKH05 and MRKH06), and four by MRKH type II (MRKH01, MRKH03, MRKH08, MRKH10) (Table 1).

We found a 1.4 Mb deletion at 17q12 in proband MRKH01. This is one of the most common chromosomal changes found in MRKH syndrome and contains the candidate genes *LHX1* and *HNF1B*. The same participant displayed also a 370 Kb interstitial duplication at 1q21.1. This is reciprocal to the TAR (thrombocytopenia, absent radius) microdeletion syndrome region, which has been reported in individuals with developmental delay and intellectual disability with incomplete penetrance and variable phenotypes (Coe 2014), although these features did not apply to our case.

The other chromosomal variants have not previously been described in MRKH syndrome, although some of them involved known candidate genes. These included *TBC1D16*, found in the 17q25 duplicated region of proband MRKH05 and reported in MRKH syndrome (Pan 2019). In addition, the 0.4 Mb duplication at 3q13.12q13.13 of proband MRKH03 contained the gene *FAM155A*, which participates in the signaling of AMH, a candidate for MRKH syndrome with unknown clinical significance (Li 2016, Mishina 1996, Oppelt 2005). Proband MRKH10 showed a microdeletion at Xp22.33 that was located within the upstream conserved noncoding elements enhancer region of *SHOX*, a gene associated with MRKH syndrome (Gervasini 2010).

Other variants had no known association with MRKH syndrome. The novel duplication at 2q24.2 involved PLA2R1, ITGB6, and RBMS1, which are all expressed in the reproductive tract. However, these genes have been associated with amelogenesis imperfecta and membranous nephropathy, and our case MRKH06 was affected by MRKH type I with no additional reported anomalies (Wang 2014, Stanescu 2011). Finally, we found long continuous stretches of homozygosity (LCSH) in two participants. Although LCSH can be associated with recessive Mendelian disorders, they usually represent ancestral haplotypes of low-recombination regions and are unlikely to be of clinical significance (Sund 2014).

### Identification of single nucleotide variants by whole exome sequencing

To provide a more complete genomic survey of our cohort, we performed WES analysis. We bioinformatically filtered the identified single nucleotide variants (SNV) against an internal database (Backhouse 2019). Following further screening for allele frequency (<0.1%), type (nonsynonymous, splice site, frameshift), and CADD score, 26 variants were selected (Table 2). All variants were heterozygous and analysis of parental DNA, where available, showed their heritability.

**Table 2.** Summary of variants identified by whole exome sequencing.

| Patient | MRKH Type | Chr | Gene | Genomic change | Protein change | Reference transcript | GnomAD allele frequency | ExAC allele frequency | CADD score | Polyphen | Relatives carrying same variant |
| --- | --- | --- | --- | --- | --- | --- | --- | --- | --- | --- | --- |
| MRKH01 | II | 15 | TLE3 | g70347561 C>T | p.Gly472Ser | NM_001105192 | 01.78E-5 | 8.36E-6 | 25.2 | PD | - |
| MRKH02 | II | 13 | FREM2 | g39338479 T>C | p.Trp1768Arg | NM_207361.6 | 0 | 0 | 28.8 | PD | Mother |
| MRKH03 | II | 5<br>16 | PROP1<br>WWOX | g177420042 A>T<br>g79245626 C>T | p.Phe117Ile<br>p.Thr393Met | NM_006261.4<br>NM_016373.4 | 0.1.08E-4<br>5.20E-4 | 1.47E-4<br>4.48E-4 | 25.1<br>19.6 | PD<br>LB | Mother<br>* |
| MRKH04 | II | 16<br>3<br>17<br>11 | TBX6<br>WNT7A<br>TBX2<br>DBX1 | g30102151 C>A<br>g13896294 C>T<br>g59479113 T>C<br>g20181818 C>T | p.Ser94Ile<br>p.Arg102Gln<br>p.Met155Thr<br>p.Arg18Leu | NM_004608.4<br>NM_004625.4<br>NM_005994.4<br>NM_001029865.4 | 0<br>2.43E-5<br>8.87E-5<br>0 | 0<br>3.34E-5<br>6.68E-5<br>0 | 23.1<br>26.9<br>28.8<br>31.0 | PD<br>PD<br>PD<br>PD | Mother, brother<br>Mother, brother<br>Mother, brother<br>Father |
| MRKH05 | I | 2<br>2<br>3 | NBAS<br>EVX2<br>FLNB | g15448404 T>C<br>g176944951 A>G<br>g58111467 C>G | p.Tyr1578Cys<br>p.Cys439Arg<br>p.Thr1353Ser | NM_015909.4<br>NM_001080458.1<br>NM_001164317.2 | 3.58E-5<br>6.70E-5<br>5.73E-4 | 3.30E-5<br>0<br>5.62E-4 | 26.6<br>25.2<br>27.7 | PD<br>PD<br>PD | - |
| MRKH06 | I | 4<br>X | FRAS1<br>DACH2 | g79458320 C>T<br>g86067976 C>T | p.Pro3755Leu<br>p.Ser453Phe | NM_025074.7<br>NM_053281.3 | 9.99E-4<br>0 | 9.55E-4<br>0 | 32.0<br>25.3 | PD<br>PD | - |
| MRKH07 | - | 19 | AMH | g2251137 C>G | p.Asp288Glu | NM_000479.5 | 1.27E-3 | 1.31E-3 | 23.5 | PD | - |
| MRKH08 | II | 22 | BCR | g23653975 T>TCCGG | p.Val1094fs | NM_004327.4 | 0 | 0 | NA | - | - |
| MRKH09 | I | 19<br>5 | B9D2<br>MAP3K1 | g41860624 C>T<br>g56160660 A>T | p.Arg170His<br>p.Met312Leu | NM_030578.4<br>NM_005921.2 | 1.06E-5<br>5.61E-5 | 1.66E-5<br>7.45E-5 | 24.8<br>23.6 | PD<br>PD | *<br>* |
| MRKH10 | II | 4<br>4 | FRAS1<br>FRAS1 | g79291045 C>T<br>g79460451 C>T | p.Arg926X<br>p.Arg3768Cys | NM_025074.7<br>NM_025074.7 | 4.02E-6<br>6.34E-5 | 0<br>3.31E-5 | 40.0<br>32.0 | NA<br>PD | - |
| MRKH11 | - | 13<br>5<br>16 | PDX1<br>NKX2-5<br>MMP2 | g28498711 C>T<br>g172660374 C>T<br>g55530864 G>A | p.Pro242Leu<br>p.Arg143Gln<br>p.Arg500His | NM_000209.4<br>NM_001166176.2<br>NM_004530.6 | 1.22E-3<br>1.25E-3<br>1.38E-3 | 2.96E-3<br>8.17E-4<br>1.41E-3 | 16.55<br>NA<br>21.6 | LB<br>-<br>PD | - |
| MRKH12 | II | X<br>14 | KLHL4<br>ESR2 | g8687731 C>G<br>g64701824 G>A | p.Leu349Val<br>p.Arg424Trp | NM_019117.5<br>NM_001437.3 | 6.01E-5<br>3.19E-5 | 6.85E-5<br>3.31E-5 | 18.76<br>24.7 | PD<br>PD | - |
| MRKH13 | I | X<br>8 | DACH2<br>ZFPM2 | g85403800 G>A<br>g106431420 A>G | p.Gly59Asp<br>p.Glu30Gly | NM_053281.3<br>NM_012082.4 | 5.46E-4<br>2.69E-3 | 6.08E-4<br>2.71E-3 | 22.3<br>25.1 | PD<br>PD | Mother<br>Father |

PD = probably damaging; LB = likely benign; NA = not available; \* = DNA available from only one parent, who did not carry the variant.

Some of the identified candidates have been linked to conditions associated with Müllerian anomalies. These included *FRAS1* and *FREM2*, both associated with Fraser syndrome, a condition characterized by cryptophthalmos, genital malformations, syndactyly, and renal anomalies (Saisawat 2012). In addition, a variant of the *NBAS* gene was found in participant MRKH05. This gene is associated with short stature optic atrophy Pelger Huët anomaly (SOPH) syndrome, in which a hypoplastic uterus is part of the clinical presentation (Maksimova 2010).

Several variants involved candidate genes that have been reported in MRKH syndrome or are known to regulate mammalian MD development. These included *TBX6*, *WNT7A*, *DACH2*, and *AMH*. *TBX6* is localized in 16p11.2 and copy number variants of this region are associated with MRKH syndrome (OMIM #611913) (Nik-Zainal 2011; Sandbacka 2013, Pontecorvi 2021). In addition, *TBX6* single nucleotide variants have been found in women affected by MRKH syndrome (Mikhael 2021). *Wnt7a*, *Dach2*, and *Amh* are involved in the formation, patterning, and development of the Müllerian ducts but their role in MRKH syndrome is not clear (Mishina 1996, Kyei-Barffour 2021). In addition, we identified variants in *EVX2*, *DBX1*, *MAP3K1*, *ESR2*, and *ZFPM2*, which play critical roles in the development of the reproductive system but are novel candidates for MRKH syndrome.

These analyses further confirm the heterogeneity of genomic findings, characterized by the identification of several variants associated with MRKH syndrome and possibly reflecting the variability in phenotypes. However, confirmation of their clinical significance requires additional functional investigation.

### In vivo functional validation of candidate gene HNF1B

The deletion at 17q12 is well known for being associated with MRKH syndrome. In our cohort, we found it in 1/13 cases (7%), in accord with the reported prevalence of 2-9% (Sandbacka 2013, Ledig 2011, Nik-Zainal 2011, Morcel 2012, Cheroki 2008, Edghill 2006). This copy number variant encompasses both *LHX1* and *HNF1B* genes, but only the role of *Lhx1* in female reproductive tract development has been assessed in vivo (Kobayashi et al., 2004; Huang et al., 2014).

To identify the specific function of *HNF1B* and its contribution to the pathogenesis of MRKH syndrome, we generated and studied a ‘conditional’ loss-of-function model in mice using a Cre/lox recombination system. This strategy requires a mouse line expressing a tissue specific Cre recombinase and a line harboring a gene of interest flanked (“floxed”) by loxP sequences. When the two lines are crossed, the Cre protein recombines the loxP sites only in the tissue where it is expressed causing deletion of the gene of interest. To ablate *Hnf1b* specifically in the epithelial cells of the MD, we crossed a *Wnt7a*-Cre line (Heliot et al., 2013; Huang et al., 2014) with a floxed *Hnf1b* mouse line (*Hnf1b^fl/fl^).* We first used the ROSA26-LacZ mouse reporter line to verify the expression of the Cre recombinase in the *Wnt7a*-Cre mouse line. We found that Cre was expressed throughout the entire period of Müllerian duct formation, as expected (Huang et al. 2014). At 11.5 days post coitum (dpc), a few mesonephric epithelial cells begin differentiation into Müllerian duct epithelial cells (Suppl Figure 3A). This is followed by the formation of the tubular structure of the Müllerian ducts, which elongate posteriorly until they reach the urogenital sinus at 13.5 dpc (Suppl Figure 3A). As demonstrated by β-gal activity, Cre expression was detected in the epithelium of the MD but not Wolffian ducts (WD) (Suppl Figure 3B).

**Figure 03.**
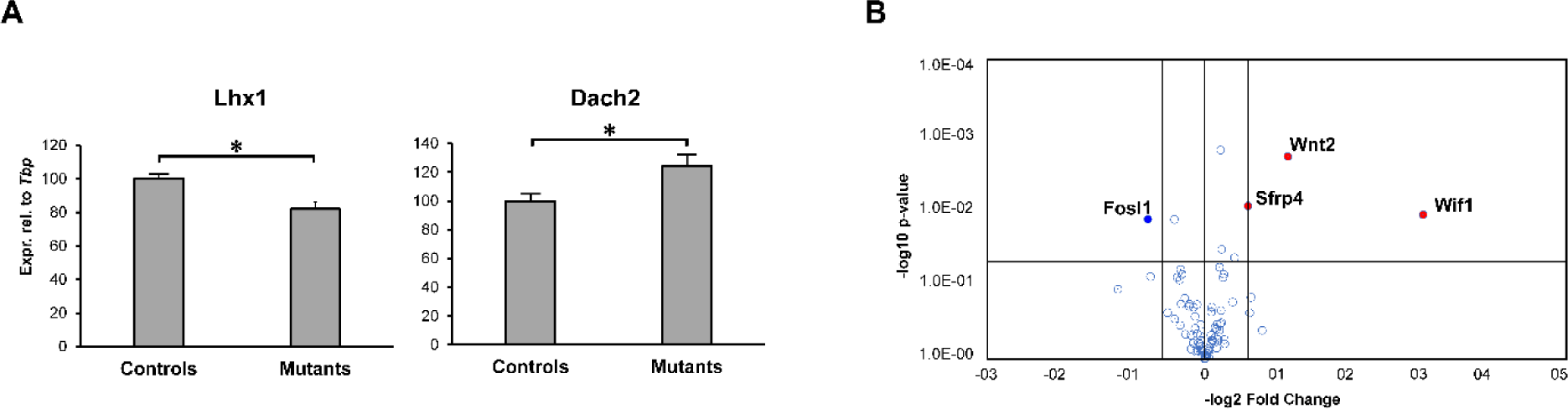
Ablation of Hnf1b cause dysregulation of few key genes involved in MD development. (A) Real-time PCR of control and Hnf1b mutant samples at 13.5 dpc showing dysregulation of Lhx1 and Dach2. (B) Volcano plot of the Wnt pathway RT2 Profiler PCR Array. Upregulated genes are in red, downregulated in blue. * = p < 0.05.

To confirm specific ablation of *Hnf1b* in *Wnt7a-Cre^+^;Hnf1b^fl/fl^* embryos, we performed immunofluorescence staining at 13.5 dpc – when both the MD and WD are normally present – and birth (P0), when only the MD remains in female animals. In non-Cre expressing littermates (*Wnt7a-Cre^-^;Hnf1b^fl/fl^* mice, referred to as controls), HNF1B protein was expressed in both the MD and WD at 13.5 dpc and in the MD at P0, as expected (Figure 1 A and B). However, the MD of *Wnt7a-Cre^+^;Hnf1b^fl/fl^* mice (referred to as mutants) were negative for HNF1B at both stages (Figure 1 A and B). We confirmed these results by qRT-PCR analysis, showing a reduction of *Hnf1b* mRNA levels by 40% compared to controls, consistent with loss of expression in the MD but not WD (Figure 1C). At P0, however, when the WD has regressed, *Hnf1b* expression was abolished in mutants (Figure 1C). These results indicate successful ablation of *Hnf1b* in the developing MD of mutant animals.

### Conditional ablation of Hnf1b results in uterine hypoplasia and renal anomalies

We examined the development of the female reproductive tract in *Wnt7a-Cre^+^;Hnf1b^fl/fl^* embryos to determine whether deletion of *Hnf1b* in mice has any consequences that mirror the phenotype of MRKH women. At embryonic stages, there were no gross differences between mutant and control MDs. However, by P0, the uteri of mutant mice appeared smaller, less developed, and were ∼20% shorter than controls than controls (Figure 2 A-C). Hematoxylin and eosin staining revealed that the mutant epithelium failed to organize in the typical pseudostratified columnar fashion of controls (Figure 2D). At P0, the stromal compartment showed reduced cell density and overall, cross-sections of the mutant uterus were smaller compared to controls (Figure 2D).

In adult mutant mice at 4 months of age, the uterus remained smaller, characterized by a thin epithelium, a stroma lacking endometrial glands, and a thicker outer longitudinal myometrium (Figure 2D). Despite the uterine hypoplasia, and similar to MRKH syndrome, the ovaries of mutant animals did not show any difference compared to controls, displaying follicles at every stage of folliculogenesis. Consistent with normal follicle dynamics, mutant female mice mated without apparent problems as evidenced by the presence of regular plugs. As expected, however, they failed to remain pregnant (Suppl Fig 4).

**Figure 04.**
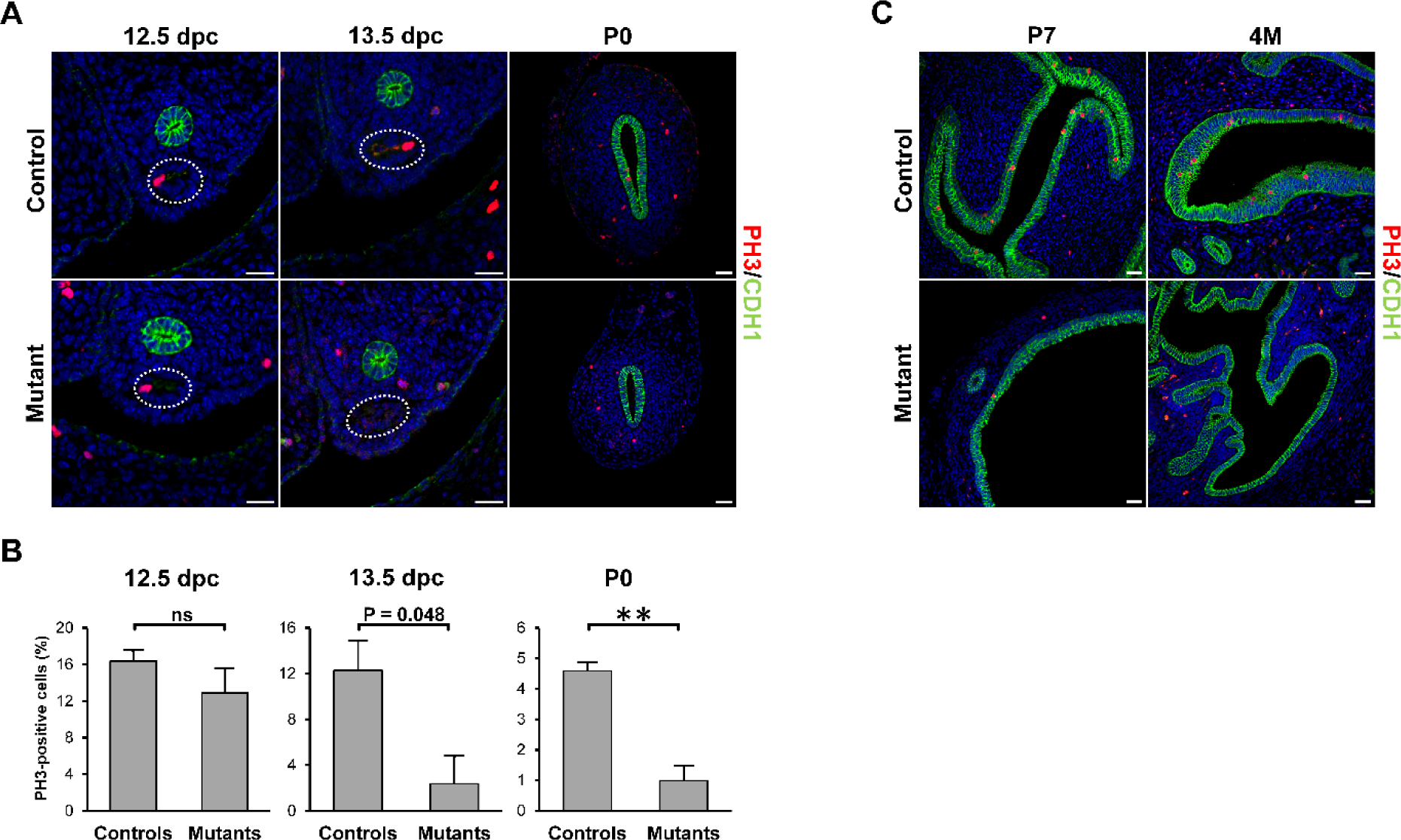
Decrease in cell proliferation in Hnf1b mutants. (A,C) Immunofluorescence staining of the proliferation marker PH3 in samples at 12.5 and 13.5 dpc, and P0, P7, and 4M. (B) Quantification of PH3-positive cells in 12.5-, 13.5 dpc, and P0 samples. Scale bar = 25 μm. * = p < 0.05, ** = p < 0.01.

Surprisingly, we found that the uterine phenotype was associated with renal anomalies similar to those described for MRKH. We detected unilateral kidney agenesis in 6/36 mutant mice (17.1%), whereas it was never observed in controls (n = 35, p = 0.0246) or heterozygous animals (n = 64, p = 0.0016) (Figure 2E). In addition to agenesis, we found cystic kidneys as additional renal association (Figure 2F), but did not assess their frequency due to the more subtle phenotype, which could not be identified a priori. These results show that *Hnf1b* ablation results in uterine hypoplasia associated with kidney anomalies, providing a mouse model for MRKH syndrome type II.

### Hnf1b regulates female reproductive tract differentiation

To investigate the molecular mechanisms by which the reproductive tract and renal anomalies in *Wnt7a-Cre^+^;Hnf1b^fl/fl^* mice arise, we next examined the expression of several known markers of MD development, beginning at 13.5 dpc. At this stage, we found a significant dysregulation of the genes *Lhx1* and *Dach2* (Figure 3A). However, several other factors including *Wnt4*, *Wnt5a*, *Wnt7*, *Wnt9*, *Dach1*, *Hoxa11*, and *Pax2* were not differentially expressed between mutants and controls (Suppl Fig 5A).

**Figure 05.**
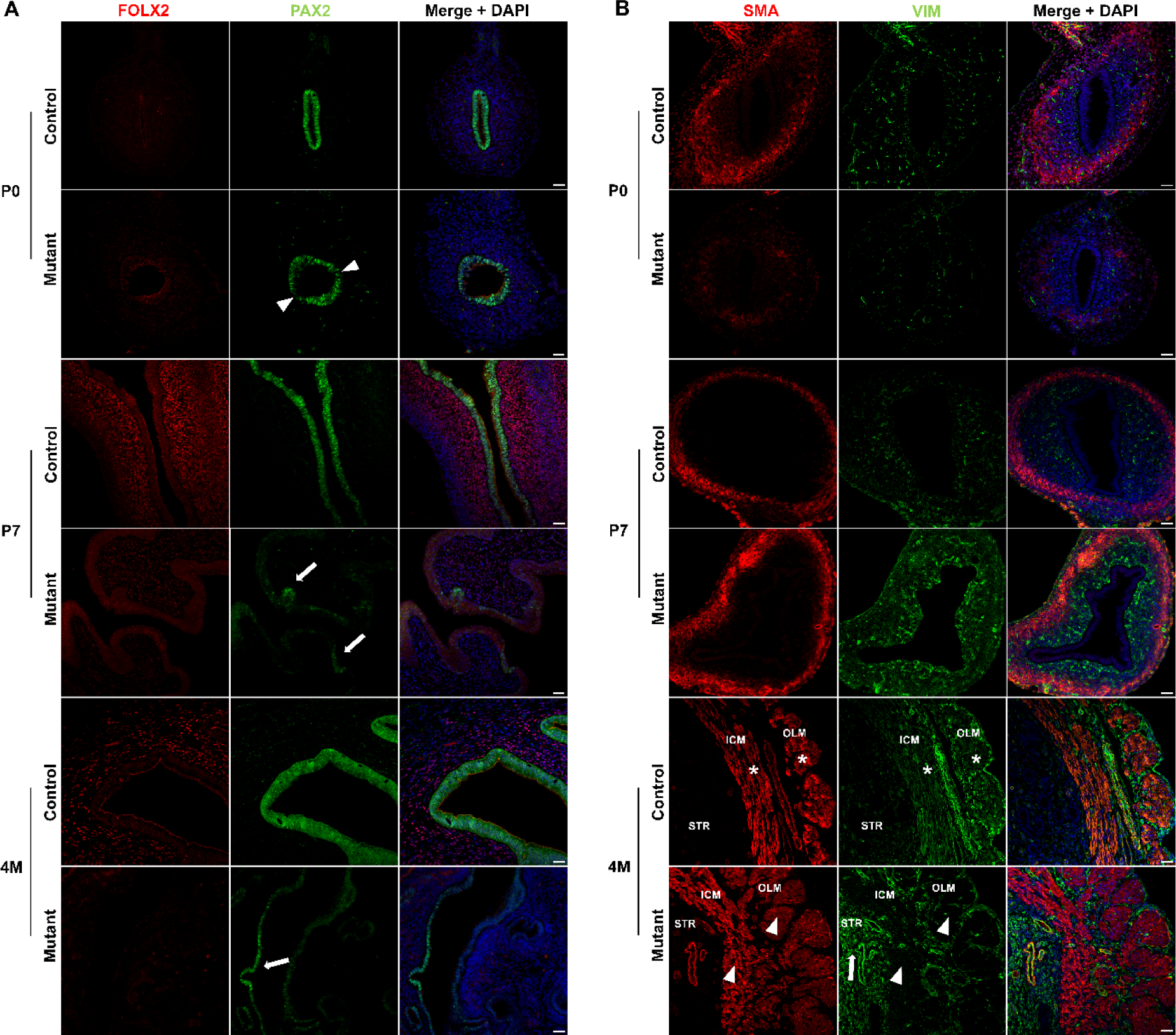
Dysregulation of epithelial and stromal markers in Hnf1b mutant uteri. (A) Immunofluorescence staining of FOXL2 and PAX2 in samples at P0, P7, and 4M. After birth, FOXL2 failed to be upregulated whereas PAX2 lost expression in mutant samples. (B) Immunofluorescence staining of SMA and VIM in samples at P0, P7, and 4M. In mutants, SMA and VIM had reduced expression at P0, but were upregulated at P7. By 4M, SMA showed the development of a thicker myometrium and VIM continued to be upregulated in the stroma (arrow) of mutant animals. VIM expression co-localized with SMA in both the inner circular (ICM) and outer longitudinal myometrium (OLM) of control (asterisks) but not mutant (arrowheads) samples. Scale bar = 25 μm.

Because of the importance of the Wnt pathway in tissue morphogenesis, we further analyzed Wnt gene expression using an RT2 Profiler PCR Array. The upregulated genes were *Wnt2*, and the Wnt-inhibitors *Wif1* and *Sfrp4*, whereas the Wnt target *Fosl1* was downregulated (Figure 3B). As Wnt signaling plays critical roles in cell proliferation, we performed immunofluorescence staining of the MD using PH3 as proliferation marker. At 12.5 dpc, there was no difference in MD epithelial cell proliferation between mutants and controls (Figure 4A,B). However, at 13.5 dpc, proliferation started to decrease in the mutants compared to controls (p = 0.048) and was strikingly reduced by P0 (Figure 4A,B), which marks the start of the differentiation of the epithelial, stromal, and smooth muscle compartments. These results are consistent with the reduced proliferation of the MD epithelium in Hnf1b mutant mice, resulting in the observed shorter uterine length. Furthermore, epithelial cell proliferation was almost completely absent in young (P7) and adult (4-month-old, 4M) uterine tissues (Figure 4C), similar to histologic observations of uterine rudiments in women with MRKH syndrome (Rall 2013).

We further characterized the uterine hypoplasia in *Hnf1b* mutants by analyzing the expression of specific markers, including the epithelial PAX2, the stromal FOXL2 and VIM, and smooth muscle actin (SMA) from P0 to 4M. In controls, PAX2 remained highly expressed throughout development and complete maturation of the uterus. However, immunostaining of PAX2 in P0 mutant did not appear uniform (Figure 5A, arrowheads), and by P7, loss of PAX2 expression was evident and limited to few, sparse clusters of epithelial cells (Figure 5A, arrows). Similarly, at 4 months, PAX2 was expressed only in some cells of the uterine epithelium (Figure 5A, arrows). At P0, FOXL2 was not expressed in control or mutant mice. However, while it was expressed as early as P7 in controls, it failed to be properly turned on in the mutant uterus, where it was virtually undetectable by 4 months (Figure 5A). Another stromal marker, VIM, was downregulated in the mutants at P0, whereas at both P7 and 4M, it was up-regulated (Figure 5B). Similarly, SMA was also downregulated at P0, consistently with the underdevelopment of the mutant uterus compared to control (Figure 5B). At P7, however, both SMA and VIM were upregulated in mutant samples (Figure 5B). By 4M, SMA staining showed the formation of a thicker but less organized myometrium, especially in its inner circular layer, which lacked a longitudinal pattern of expression (Figure 5B). Interestingly, vimentin remained upregulated in the stromal endometrium, but showed dysregulation in the myometrium (Figure 5B, arrow). In the control uterus, VIM co-localized with the SMA in the inner circular and outer longitudinal myometrium (Figure 5B, asterisks). In the mutant, however, both myometrial layers lost expression of VIM (Figure 5B, arrowheads).

We also checked the expression of laminin, an important component of the basal lamina of epithelial and endothelial layers (Figure 6). Laminin showed high and uniform expression in control mice. In the mutants at P0, laminin showed a lower expression at the base of the MD epithelium, while remaining normally expressed around blood vessels (Figure 6 arrowheads).

**Figure 06.**
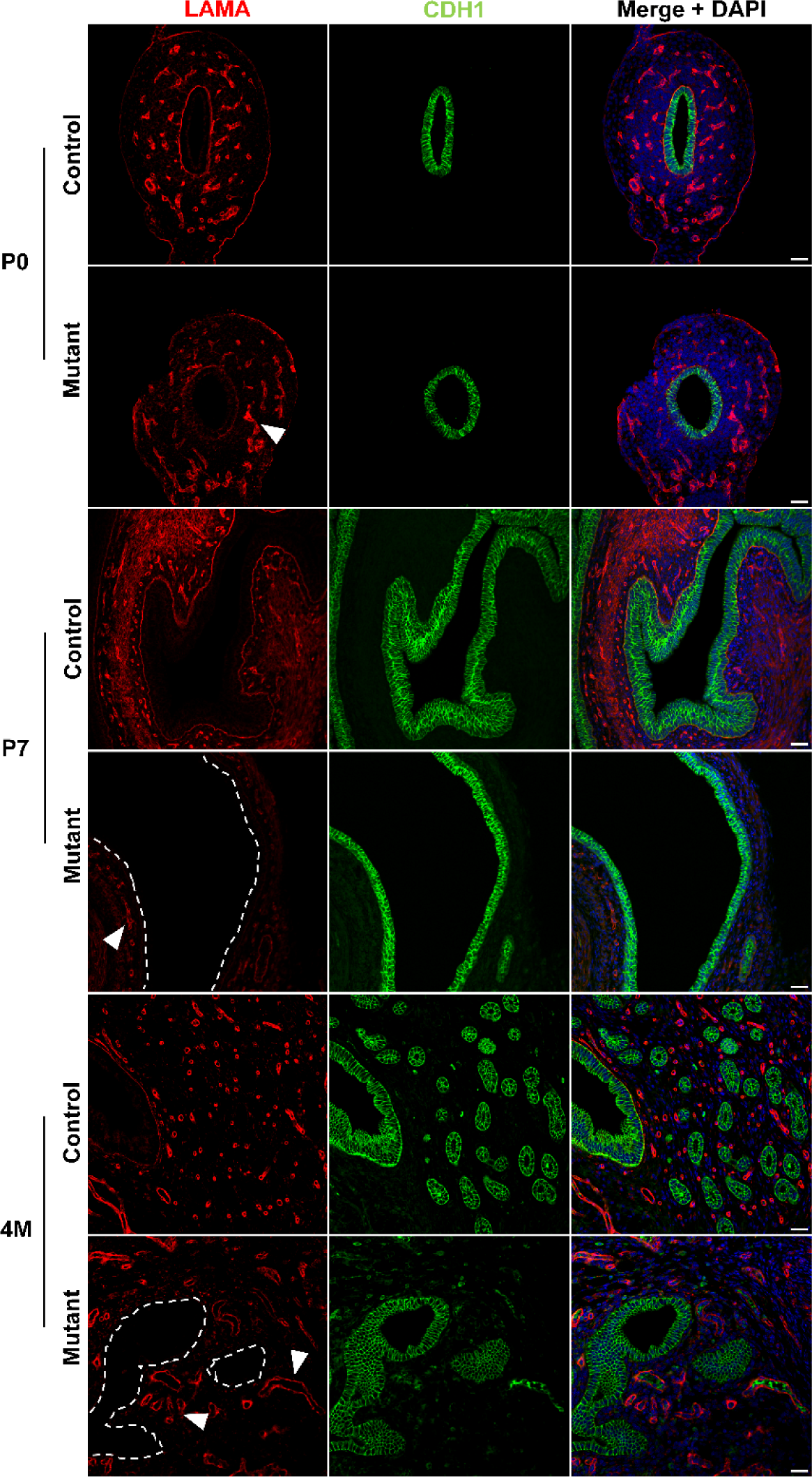
Laminin is lost in the uterine epithelium basal lamina. Immunofluorescence staining of LAMA and CDH1 in control and mutant samples at P0, P7, and 4M. Laminin is gradually lost underneath the uterine epithelium (dashed lines represent the location of the missing LAMA staining). In mutant samples, LAMA was expressed around the vasculature (arrowheads) similar to controls. Scale bar = 25 μm

By P7, and showing also in 4M samples, the mouse uterus had completely lost expression of laminin underneath the MD epithelium (Figure 6, dashed lines).

Together, these results show that key epithelial, stromal, and smooth muscle markers are dysregulated as early as P0, consistent with the uterine hypoplasia observed in mutant mice.

### Single cell transcriptomics of the differentiating uterus

Although *Hnf1b* was ablated specifically in the MD epithelium, uterine hypoplasia extended to the stromal and muscular compartments of the uterus. To better understand the specific molecular and cellular changes in each cell type, we performed single cell RNA-Seq analysis at P0, the time when immature MD cells start differentiating into specialized uterine cells. Single cell transcriptomic analysis identified nine distinct cell populations in both control and *Hnf1b* mutants (Figure 7A and Suppl Figure 2). We assigned identities to these clusters based on expression enrichment of specific markers and the Mouse Cell Atlas single cell RNA-Seq database (Han 2018).

**Figure 07.**
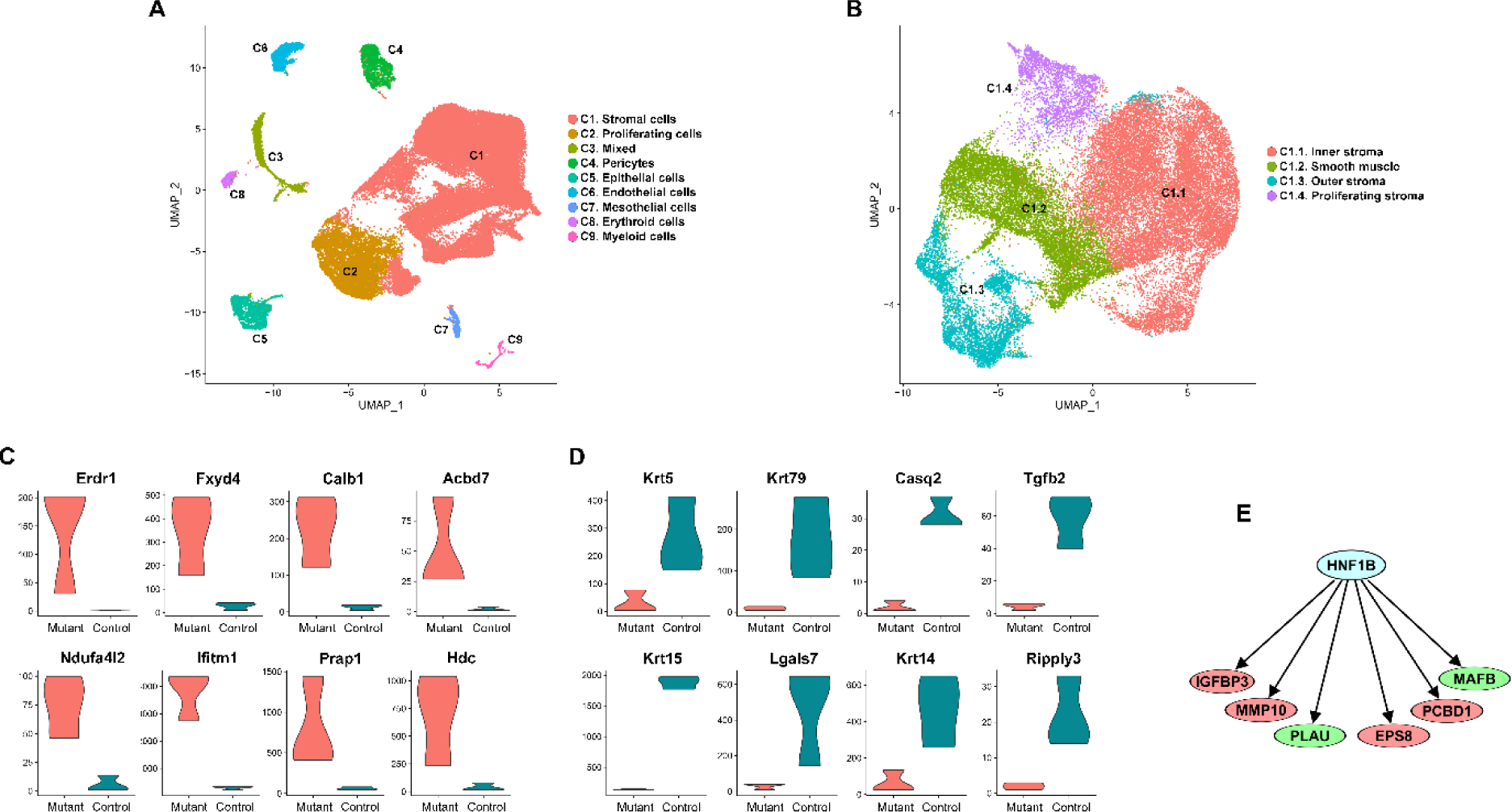
Cell population of the newborn uterus. (A) UMAP plot from sc-RNA Seq analysis of uterine samples at P0 reveals 9 cell clusters (C1- C9 from largest to smallest). (B) UMAP plot of the C1 stromal population after sub-clustering. (C) Violin plots of the top upregulated and (D) downregulated genes in the epithelial cell cluster. (E) Factors annotated as downstream of Hnf1b in the Ingenuity Knowledge Database that were differentially expressed in the Hnf1b mutant samples (red = upregulated, green downregulated).

The cluster C1 represented the most abundant cell population expressing stromal markers such as *Des*, *Htra3*, *Cnrip1*, and *Csrp1*. Cluster C2 expressed genes that are characteristic of proliferating cells, including *Stmn1*, *Top2a*, *Mki67*, *Prc1*, and *Kif11*. Cluster C3 showed genes with shared expression among several cell types, and likely represented a mixed population, or an unidentified cell type. C4 represented pericytes, which expressed the markers *Cspg4*, *Abcc9*, *Rgs5*, and *Cald1*. The cluster C5 consisted of epithelial cells, which expressed markers including *Cdh1* and *Cldn4*. Cluster C6 showed enriched expression of genes associated with endothelial cell, including *Cd93*, *Tie1*, and *Esam*. C7 expressed genes characteristic of the mesothelium such as *Msln*, *Dpp4*, *Muc16*, and *Fbln2*. Clusters C8 represented the erythroid lineage, expressing markers *Hba.a1*, *Hba.a2*, *Hbb*, and *Alas2*. Finally, C9 was made of myeloid cells, which expressed *Lyz2*, *Cybb*, and *Aif1*.

To characterize more in detail the large population of stromal cells, we performed further sub- clustering of C1, revealing four sub-types (Figure 7B). Cluster C1.1 expressed markers of the inner stroma *Vcan, Plac8*, *Cpxm2*, and *Axin2* (Saatcioglu 2019). The cluster C1.2 represented smooth muscle cells, which expressed *Ptn*, *Lum*, *Col1a1*, and *Gpc3*. The cluster C1.3 showed enrichment of outer stroma genes such as *Smoc2*, *Apoe*, and *Fbn2*. Cluster C1.4 expressed genes associated with cell cycle, including *Hmgb2*, *Tubb5*, *Cenpf*, and *Tmpo*, and represented proliferating stromal cells.

### Single cell transcriptomic profile reveals dysregulation of key pathways during reproductive tract differentiation

To gain further insight into the molecular processes that were disrupted by *Hnf1b* ablation, we analyzed the gene expression profiles of the three main clusters of epithelial, stromal and proliferating cells. Consistent with our histological findings, both Gene Ontology and Ingenuity Pathway Analysis (IPA) showed that the most affected processes involved differentiation, proliferation, and cell development (Table 3, Suppl Table 2). In addition, IPA revealed that organismal development, survival, embryonic development, and organ morphology were the most enriched functional annotations for the differentially expressed genes in our datasets (Table 4).

**Table 3.** Gene Ontology analysis of differentially expressed genes. Top 10 Biological Process classes in epithelial, stromal, and proliferating cell clusters.

|  | **Epithelial cells** |  | **Stromal cells** |  | **Proliferating cells** |  |
| --- | --- | --- | --- | --- | --- | --- |
| **Rank** | **GO class** | **# molecules** | **GO class** | **# molecules** | **GO class** | **# molecules** |
| 1 | Cell differentiation | 30 | Multicellular organism development | 20 | Multicellular organism development | 21 |
| 2 | Negative regulation of cell proliferation | 25 | Cell differentiation | 18 | Cell adhesion | 16 |
| 3 | Negative regulation of apoptotic process | 24 | Positive regulation of gene expression | 12 | Cell differentiation | 13 |
| 4 | Positive regulation of transcription | 24 | Phosphorylation | 12 | Response to hypoxia | 10 |
| 5 | Cell adhesion | 22 | Positive regulation of apoptotic process | 10 | Phosphorylation | 10 |
| 6 | Phosphorylation | 21 | Angiogenesis | 9 | Cellular response to hypoxia | 9 |
| 7 | Lipid metabolic process | 20 | Transmembrane transport | 9 | Angiogenesis | 9 |
| 8 | Positive regulation of gene expression | 20 | Nervous system development | 9 | Positive regulation of apoptotic process | 7 |
| 9 | Positive regulation of apoptotic process | 19 | Glycolytic process | 8 | Glycolytic process | 6 |
| 10 | Angiogenesis | 17 | Response to hypoxia | 8 | Negative regulation of cell migration | 6 |

**Table 4.** Pathway analysis of differentially expressed genes in epithelial, stromal, and proliferating cell clusters. Most enriched “System Development and Function” categories.

| **Epithelial cells** |  |  | **Stromal cells** |  |  | **Proliferating cells** |  |  |
| --- | --- | --- | --- | --- | --- | --- | --- | --- |
| **Category** | **No. of molecules** | **p-value range** | **Category** | **No. of molecules** | **p-value range** | **Category** | **No. of molecules** | **p-value range** |
| Organismal development | 226 | 1.72E-05 - 1.57E-14 | Organismal development | 80 | 4.29E-03 - 2.16E-05 | Organismal development | 70 | 6.68E-03 - 2.66E-05 |
| Organismal Survival | 162 | 9.42E-06 - 1.75E-16 | Organismal Survival | 66 | 5.51E-07 - 5.51E-07 | Cardiovascular System Development and Function | 36 | 6.68E-03 - 2.66E-05 |
| Embryonic Development | 127 | 1.02E-05 - 4.67E-12 | Organ Morphology | 51 | 4.50E-03 - 2.16E-05 | Connective System Development and Function | 35 | 6.68E-03 - 2.64E-05 |
| Cardiovascular System Development and Function | 117 | 1.86E-05 - 3.58E-13 | Cardiovascular System Development and Function | 47 | 3.96E-03 - 1.08E-05 | Tissue Morphology | 28 | 6.68E-03 - 2.64E-05 |
| Digestive System Development and Function | 70 | 1.33E-05 - 9.99E-12 | Reproductive System Development and Function | 28 | 4.06E-03 - 9.78E-06 | Skeletal and Muscular System Development and Function | 23 | 6.68E-03 - 8.29E-07 |

The role of *Hnf1b* in the MD epithelial cells including what genes are downstream of Hnf1b are unknown. To address this issue, we focused on the epithelial cluster to identify factors that were affected by *Hnf1b* ablation. The most strongly up- and down-regulated genes, shown in Figure 7C-D, play critical roles in several developmental processes. These factors included *Erdr1*, a negative regulator of cell proliferation and migration; *Ifitm1*, a patterning gene; and *Prap1* and *Hdc*, which are involved in cell migration. Another upregulated factor was *Ndufa4l2*, which is involved in the estrogen receptor pathway, whereas *Calb1* plays a role in kidney development. Among the most down-regulated genes, we found factors regulating epithelial cell development, including *Krt5*, *Krt14*, *Krt15*, and *Krt79*, as well as genes involved in patterning, proliferation, and migration such as *Tgf2b*, *Lgals7*, and *Ripply3*.

We then interrogated the Ingenuity Knowledge Database for all factors that have been reported as downstream of and having a direct relationship with *Hnf1b*. We overlaid our dataset of differentially expressed genes to identify factors that could be targets of HNF1B in the MD epithelium (Figure 7E). This analysis revealed the patterning gene *Mafb*, which is directly activated by HNF1B in response to retinoic acid (Sturgeon 2011), and the estrogen pathway genes *Igfbp3* and *Pcbd1*, the latter of which is a binding partner of HNF1B (Ferrè 2014). In addition, other genes included *Mmp10*, *Eps8*, and *Plau*, which are all involved in cell proliferation, migration, and differentiation. Pathway explorer analysis in IPA showed that these factors were also involved in cell polarization, which is a critical process for epithelial function (Suppl Figure 6).

Finally, we investigated which of the main canonical pathways involved in Müllerian duct development were the most disrupted. We found that the estrogen receptor, the PI3K/AKT, the retinoic acid receptor signalling, and the epithelial-mesenchymal transition pathway had the largest number of dysregulated genes, whereas WNT, TGF-β, BMP, and FGF seemed less affected (Table 5).

**Table 5.** Number of differentially expressed genes in the pathways regulating female reproductive tract development.

| Pathway | # molecules |  |  |
| --- | --- | --- | --- |
|  | Epithelial | Stromal | Proliferating |
| Estrogen Receptor | 11 | 4 | 4 |
| PI3K/AKT | 15 | 1 | 2 |
| Retinoic Acid Receptor | 10 | 2 | 0 |
| Epithelial-Mesenchymal Transition | 8 | 5 | 4 |
| WNT | 6 | 3 | 2 |
| TGFβ | 7 | 1 | 2 |
| BMP | 3 | 3 | 4 |
| FGF | 3 | 3 | 1 |

These results show that Hnf1b regulates key processes of MD development including epithelial cell proliferation, migration, and differentiation, and that its ablation results in the disruption – across multiple cell populations – of pathways that are critical for uterine differentiation.

## Discussion

MRKH syndrome is a significant women’s reproductive health issue. This congenital condition denies affected women the possibility of conceiving and bearing children, and its almost invariably unexpected diagnosis can result in distress, confusion, depression, and shame (Carroll 2020). The situation is exacerbated by the current inability of healthcare professionals to provide a molecular diagnosis and hence an explanation for how the condition arose in any given affected woman. In this light, defining the genetic causes of MRKH syndrome is an important goal but remains an ongoing challenge in clinical genomics. Several approaches have been applied including microarray analysis and genome-wide DNA sequencing (Nodale 2014, Mikhael 2021, Pontecorvi 2021). These investigations have revealed a long list of candidate genes, but confirmation of their role in MD development and most importantly MRKH syndrome is lacking.

In this study, we combined DNA analysis of women with MRKH syndrome using SNP microarray and whole exome sequencing and the functional genomics in the mouse model to validate the role of *HNF1B*. We identified several known and novel candidates for MRKH syndrome, and we chose *HNF1B* due to the lack of information on its function. *HNF1B* is located in 17q12, whose deletion is one of the most frequent chromosomal changes associated with MRKH syndrome. The same region contains also the gene *LHX1*, which has been demonstrated to be critical for MD development (Huang 2014) leaving the role of *HNF1B* uncertain.

By specific ablation in the MD epithelium, we demonstrated that *Hnf1b* is necessary for the differentiation of the female reproductive tract. *Hnf1b* mutant mice displayed a hypoplastic uterus, characterized by a simple cuboidal epithelium and reduced stromal thickness that failed to properly develop endometrial glands. The *Hnf1b* mutant uterus resembled the histology of uterine rudiments found in MRKH syndrome, including a lower cell proliferation capacity and a simple, less differentiated epithelium (Rall 2013). *Hnf1b* loss-of-function caused a decrease in MD epithelial cell proliferation, which started as early as 13.5 dpc, leading to the development of a shorter uterus. Although a few genes were dysregulated during embryonic development, including *Lhx1,* and factors of the Wnt pathway, morphological differences between *Hnf1b* mutant and control animals became evident around the time of birth when uterine differentiation takes place (Stewart 2013).

In *Hnf1b* mutant mice, the epithelium failed to maintain expression of PAX2, which is necessary for the development of the female reproductive tract (Torres 1995). Similarly, expression of laminin was lost in the postnatal mutant uterus. Laminin is a component of the basement membrane, required for maintaining polarization of the epithelial layer (Matlin 2017). As the basal lamina is also important for the functional coordination between epithelium and the underlying stroma, loss of laminin could explain the dysregulated expression of other markers including VIM and SMA, which were downregulated at birth but upregulated at later stages. In addition, the stromal FOXL2 – which is turned on after birth – failed to be expressed, leading to a phenotype similar to that of *Pgr^Cre/+^;Foxl2^fl/fl^* mutant mice and characterized by a smaller stromal compartment and a thicker myometrial layer (Bellessort 2015). This could be due to the proposed role of FOXL2 to regulate *Wnt7a*, which in turn maintains the expression of other Wnt and Hoxa factors inducing the differentiation of the stroma into the endometrial and myometrial compartments (Bellessort 2015).

Our transcriptomic analysis revealed important insight into the molecular mechanisms responsible for uterine hypoplasia in mutant mice. The main processes affected by *Hnf1b* loss included cell proliferation and differentiation in the epithelial, stromal, and proliferative cell clusters. As expected, disruption of gene expression was greater in the epithelial population compared to the stromal and proliferative clusters, and involved genes that regulate patterning, migration, and epithelial development. The most affected pathways were the PI3K, estrogen, and retinoic acid signaling, which seem to be involved in later phases of MD development and differentiation rather than initial formation. The PI3K/AKT pathway is believed to play a role in the breaking down of the extracellular matrix to allow for MD elongation (Fujino 2009). The estrogen receptor pathway is critical for uterine and vaginal epithelial proliferation as well as uterine gland development in pubertal mice (Nanjappa 2015). Retinoic acid signaling is necessary for MD epithelial and stromal differentiation, as well as maintenance of uterine homeostasis in the adult (Nakajima 2019). In addition, we found the downregulation of critical factors including *Lef1*, *Wnt6*, and *Wnt10a*, suggestive of the inhibition of the Wnt pathway, which plays critical roles in cell differentiation and epithelial-mesenchymal transition, both necessary for the development of the reproductive tract.

Although *Hnf1b* plays important roles in kidney development (Lokmane 2010, Heliot 2013), the finding of kidney agenesis and kidney anomalies following *Hnf1b* ablation in the MD epithelium was surprising. It is unclear why this phenotype occurred only in some mutant animals. This may be due to the Cre-directed ablation of *Hnf1b* in a subset of cells of the developing kidney. Maybe the activity of the *Wnt7a* promoter is variable in some mesonephric or metanephric cells leading to inconsistent expression of Cre and therefore ablation of *Hnf1b*. The prevalence of kidney agenesis in *Hnf1b* mutant mice was similar to that of MRKH type II (Morcel 2007), and this was reminiscent of the incomplete penetrance and variable expressivity characteristic of this condition.

Our study demonstrates the benefit of modelling MRKH syndrome in the mouse following identification of candidate genes in human. Our genomic analyses detected several known and novel copy number and single nucleotide variants, providing useful clues to the genetics of MRKH syndrome. In a previous report, we identified a *FRAS1* variant associated with this condition (Backhouse 2018), and here we found *FRAS1* variants in two additional participants. This gene is part of the basal membrane, and is required to maintain epithelial cell integrity (Saisawat 2012). In addition, we found a variant in the *FRAS1*-related factor *FREM2*, which is required for proper differentiation of epithelial cells (Jadeja 2005). We identified other genes with important roles in key pathways including *ESR2*, with a proposed mechanism in MRKH syndrome (Rall 2011); *B9D2*, involved in Hedgehog signaling (Dowdle 2011, Migone 2012); and *DBX1* involved in patterning and cell migration (Pierani 2001). However, the specific role of these candidates in MRKH syndrome remains unknown without functional validation in vivo.

Although we were able to study relatively few unaffected family members, our analysis of parental DNA provided clues about the origin of the identified variants. For example, MRKH04 was affected by MRKH type II featuring spinal anomalies and unilateral kidney agenesis. Both parents and two brothers – one of which was also affected by unilateral kidney agenesis – participated in this study. We identified variants in *TBX2*, *TBX6*, and *WNT7A*, which were inherited from the mother and shared with the affected brother, whereas the *DBX1* variant was inherited from the father but not shared with the affected brother. Although useful, this information explained only part of the etiology of MRKH especially when considering the mechanisms of incomplete penetrance and variable expressivity. It is believed that complex genetic interactions may be at the root of these phenomena, and it is possible that the presence of multiple heterozygous genetic variants – each with small but synergistic effects – could be involved. This would be consistent with the polygenic etiology proposed for some cases (Morcel 2007). However, environmental factors may also play a role, and further studies are needed to understand how they could contribute – alone or by interacting with genetic factors to the variability of conditions like MRKH syndrome.

## Conclusions

A diagnosis of MRKH syndrome has a major impact on the lives of affected individuals. Our study has provided firmer evidence for the involvement of several candidate genes, identified new potential candidate genes for further study, and most importantly provided definitive evidence that *HNF1B* variants cause MRKH features. These outcomes will aid efforts to provide informative molecular diagnosis for MRKH syndrome, and improve genetic counselling options, leading to better psychological support and promoting awareness for a condition that remains enigmatic for clinicians, scientists, affected women and the community.

## Acknowledgements

We thank MRKH Australia for their support in recruiting women, Dr. Richard Behringer for providing the *Wnt7-Cre* mouse line, Dr. Katrina Bell for assistance with variant analysis, and Enya Longmuss for assistance with managing mouse colonies. This work was supported by research grants from the National Health and Medical Research Council of Australia.

