## Supplementary material for "Functional genomics analysis identifies impairment of *HNF1B* function as a cause of Mayer-Rokitansky-Küster-Hauser syndrome"

**Supplementary Figures**

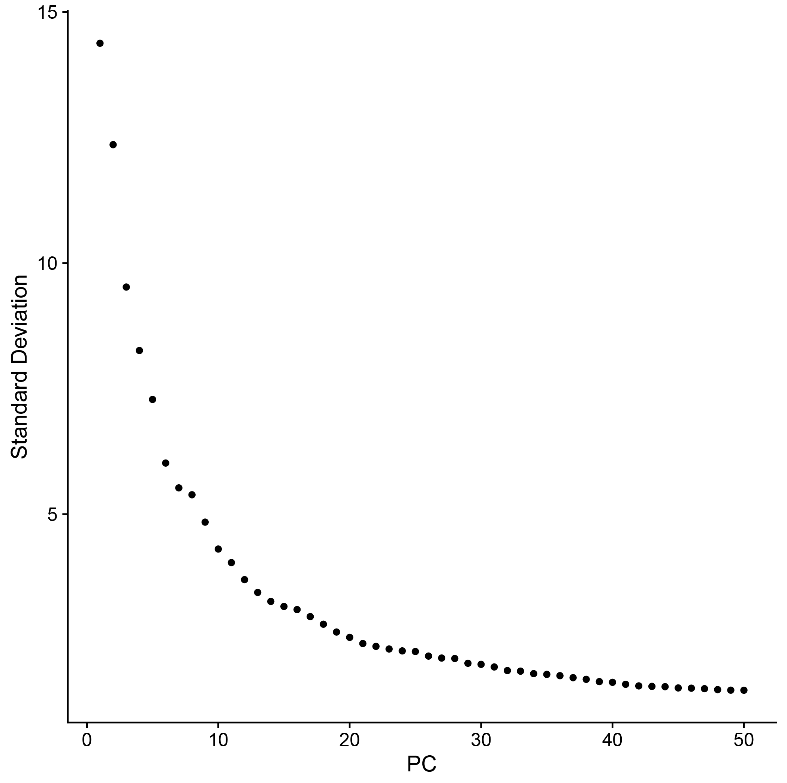

**Suppl Figure 01. Elbow plot of principal components.**

Elbow plot of the principal components (PC) ranked based on the percentage of variance explained after Principal Component Analysis.

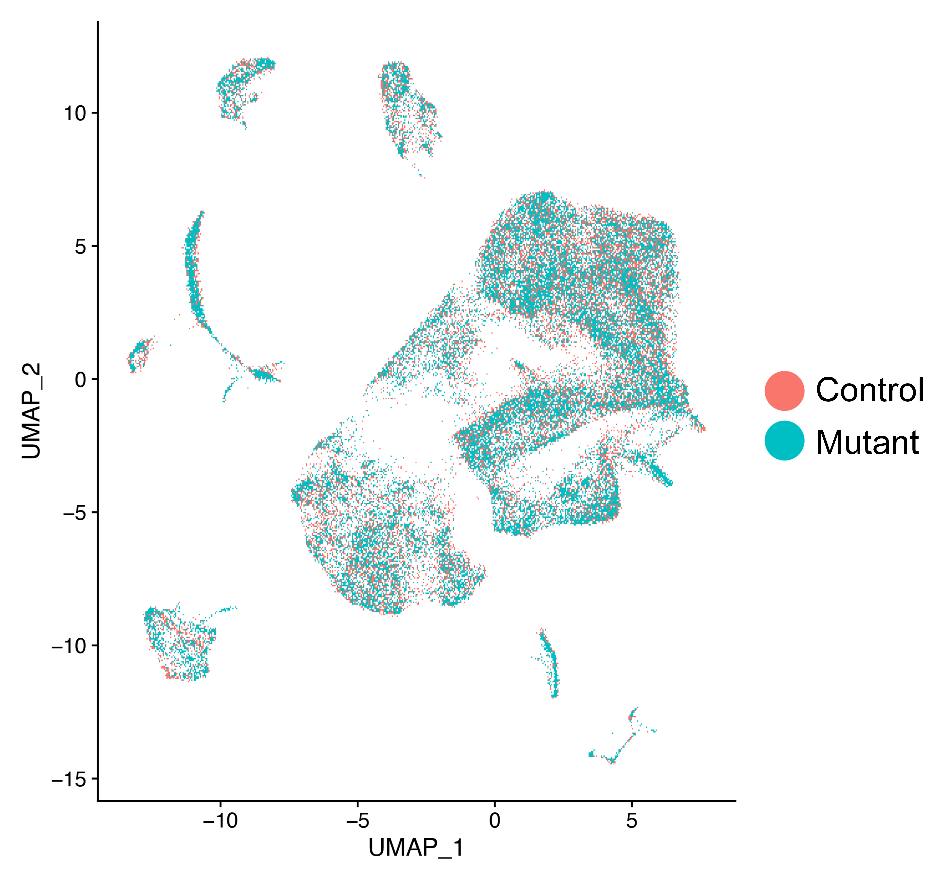

**Suppl Figure 02. Uniform manifold approximation and projection of single cell analysis.**

Uniform manifold approximation and projection (UMAP) plot shows data integration from control and Hnf1b mutants.

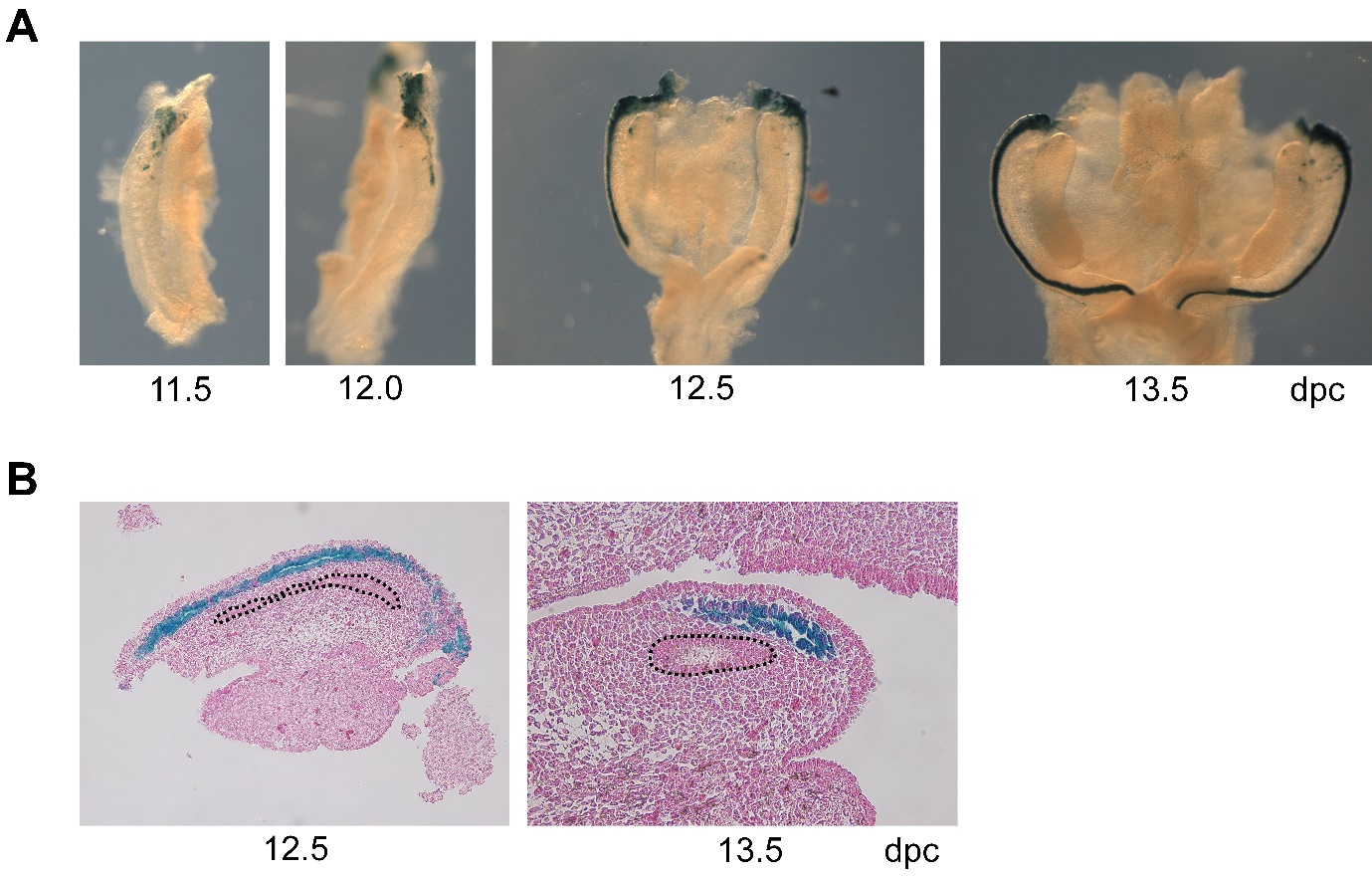

**Suppl Figure 03. Characterization of Wnt7a-Cre expression.**

(A) LacZ staining of *Wnt7a-Cre mice crossed with* ROSA26-LacZ mice showing Wnt7a-directed expression of Cre started at 11.5 dpc at the tip of the MD and progressed posteriorly during MD elongation. (B) Sections of LacZ-stained samples showed Wnt7a-Cre expression in the MD (blue) but not the WD (dashed lines). Counterstaining was performed with Nuclear Fast Red.

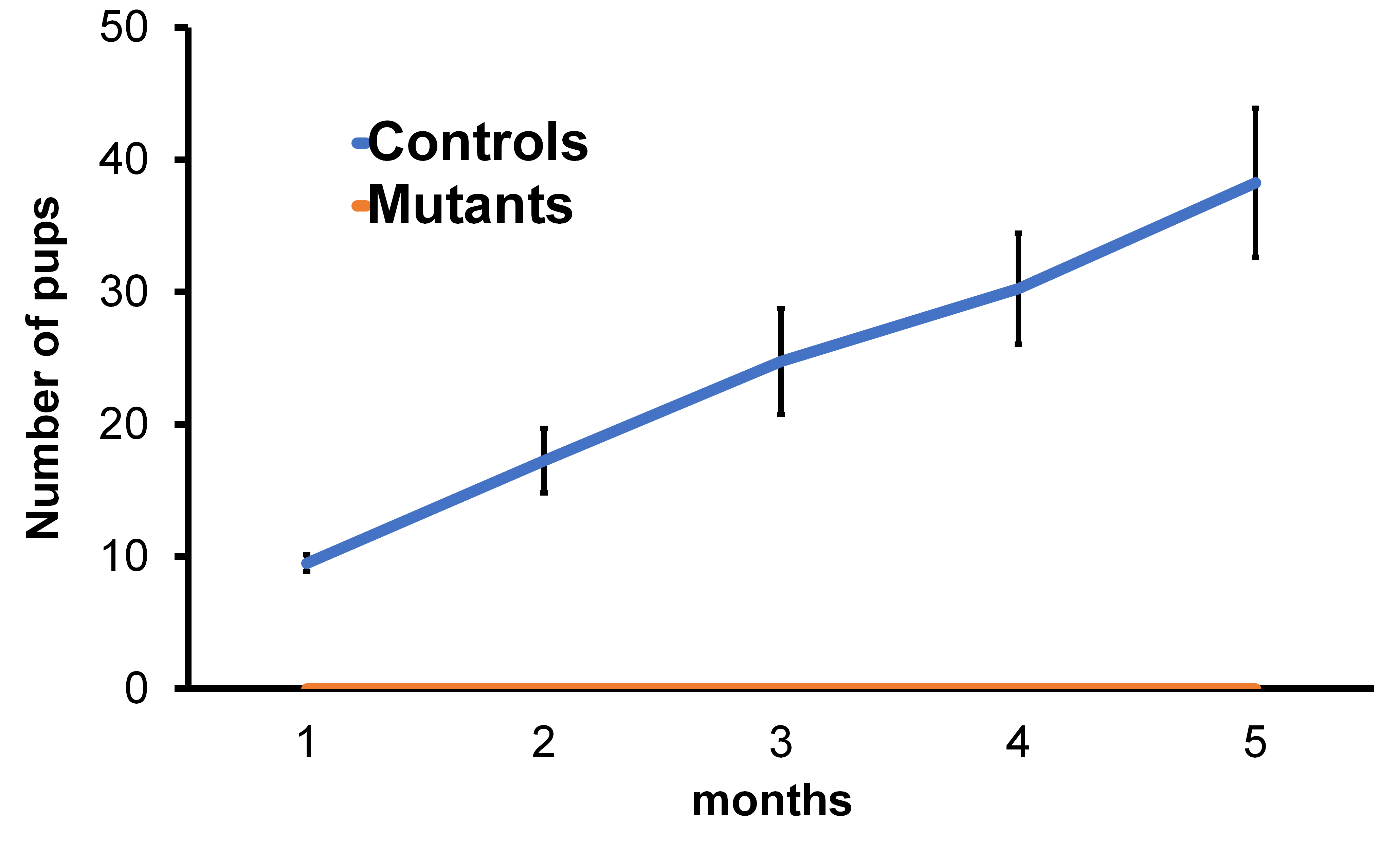

**Suppl Figure 04. Hnf1b-mutant mice fail to remain pregnant.**

Progeny analysis showed that Hnf1b mutant mice failed to remain pregnant and deliver pups.

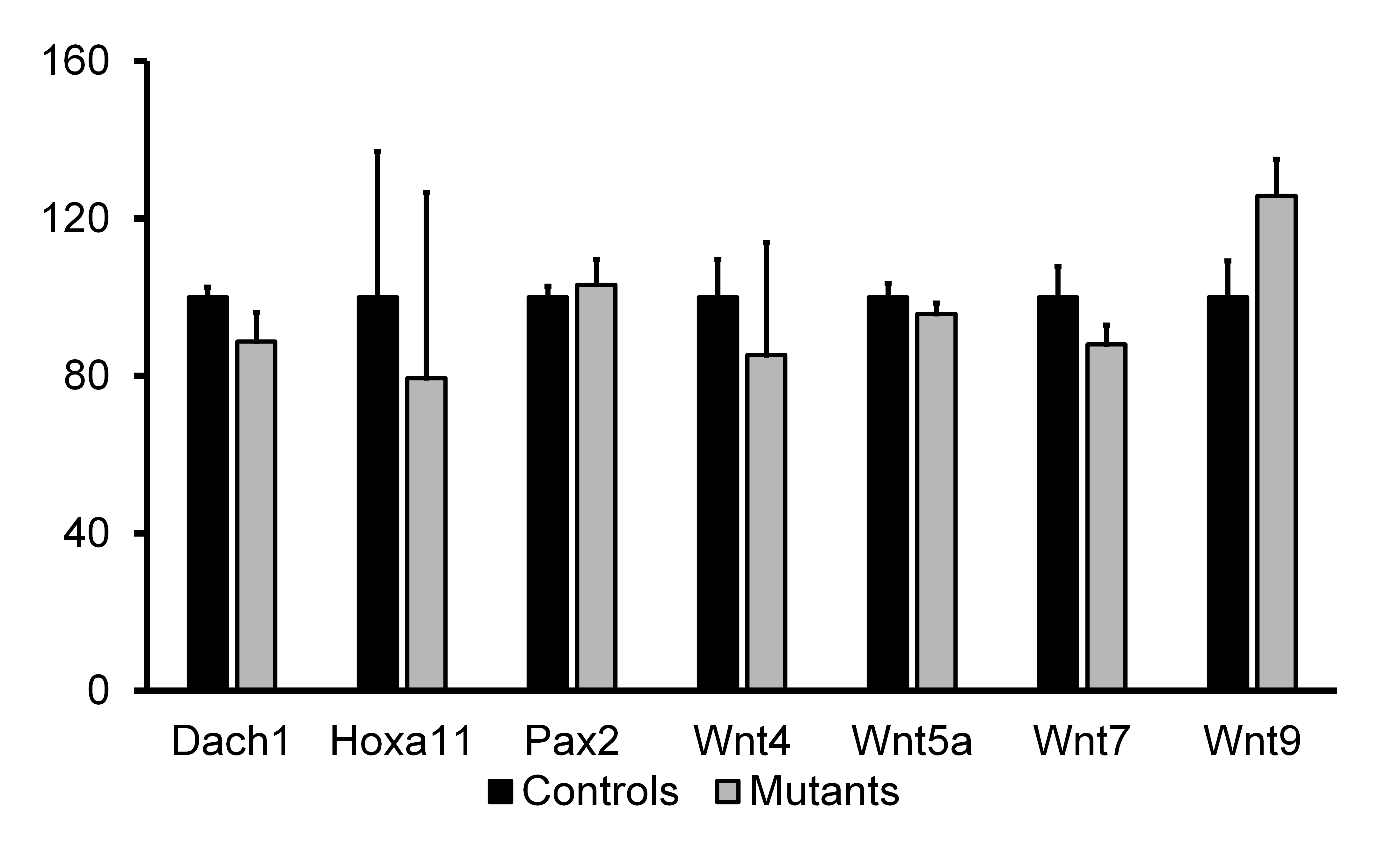

**Suppl Figure 05. Gene expression of markers of MD development in 13.5 dpc samples.**

Expression of several factors known to play an important role in MD development was not significantly different between controls and Hnf1b mutant samples.

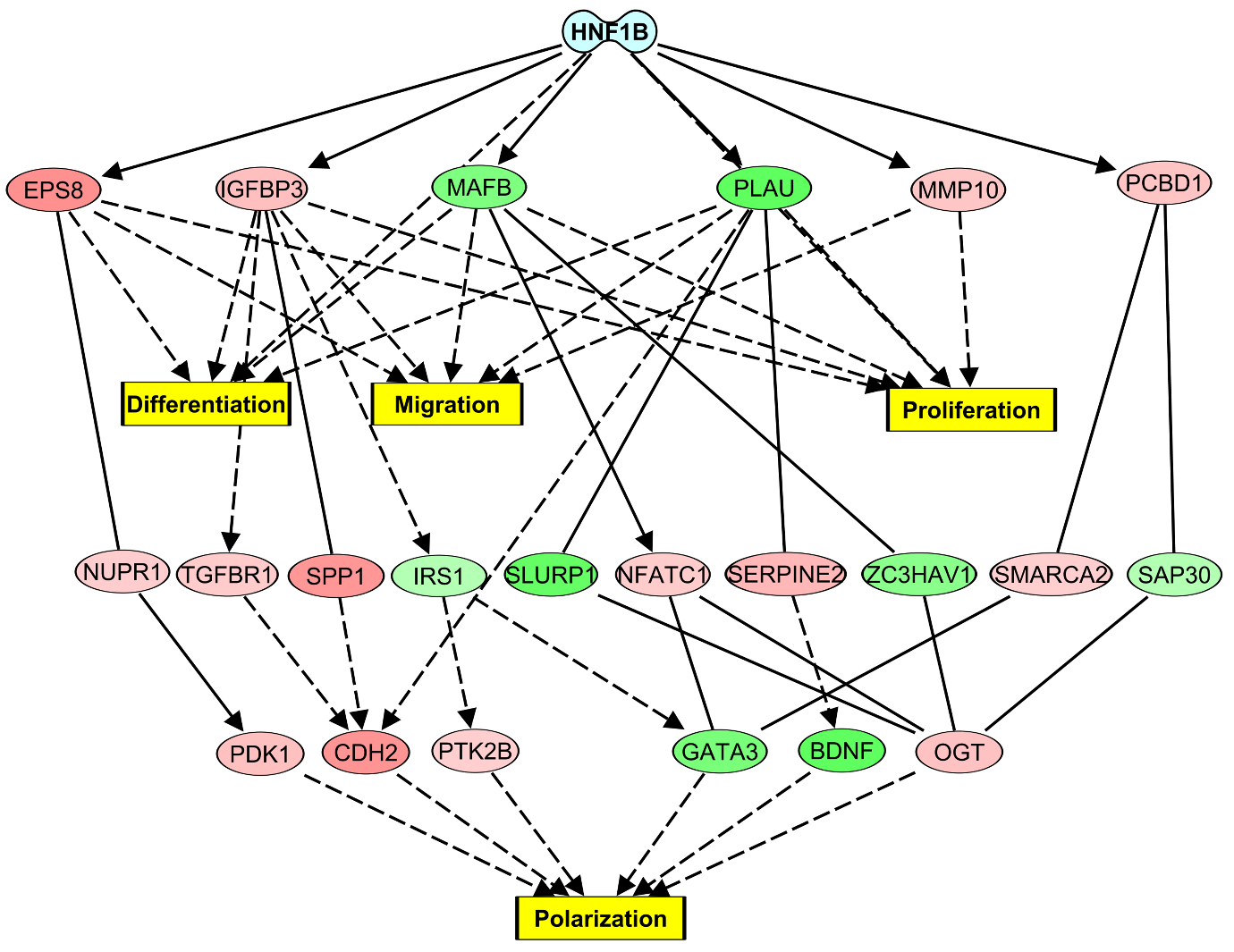

**Suppl Figure 06. Differentially expressed genes downstream of Hnf1b are involved in key processes of female reproductive tract development.**

Pathway explorer analysis of differentially expressed genes downstream of Hnf1b in the epithelial cell cluster revealed their role in cell differentiation, migration, and proliferation as well as polarization.

**Suppl Table 1. List of primers for real-time PCR**

| Hnf1b-Fw | CTATAGCTCCAACCAGACGC |
| --- | --- |
| Hnf1b-Rev | AGTGACCTCATTGTTTCCCG |
| Wnt9-Fw | GGGACAACCTCAAGTACAGC |
| Wnt9-Rev | TTCCACTCCAGCCTTTATCAC |
| Dach1-Fw | ACTTTCTCTAACTGGGCATGG |
| Dach1-Rev | AGCTCTGGCATTGTCTATGG |
| Dach2-Fw | CTCCGACCCTTAATCCACTTC |
| Dach2-Rev | ATTCATCTGATTCATTGCCATGG |
| Hoxa11-Fw | AGGAGAAGGAGCGACGG |
| Hoxa11-Rev | GGTATTTGGTATAAGGGCAGCG |
| Wnt7a-Fw | ACGAGTGTCAGTTTCAGTTCC |
| Wnt7a-Rev | AATCGCATAGGTGAAGGCAG |
| Lhx1-Fw | CCAGTGCTGTGAATGTAAATGC |
| Lhx1-Rev | GAACCAGATCGCTTGGAGAG |
| Pax2-Fw | GATCCTACTCCATCAACGGG |
| Pax2-Rev | GACCAGATGTAAACCTCCACC |
| Wnt4-Fw | CTGGACTCCCTCCCTGTCTTT |
| Wnt4-Rev | CATGCCCTTGTCACTGCAA |
| Wnt5A-Fw | AATGAAGCAGGCCGTAGGA |
| Wnt5A-Rev | AGCCAGCACGTCTTGAGG |
| TBP-Fw | ACGGACAACTGCGTTGATTTT |
| TBP-Rev | ACTTAGCTGGGAAGCCCAAC |
| Hnf1b-Fw | CTATAGCTCCAACCAGACGC |

**Suppl Table 2. Pathway analysis of differentially expressed genes in epithelial, stromal, and proliferating cell clusters. Most enriched “Molecular and Cellular Functions” categories.**

| **Epithelial cells** | | | **Stromal cells** | | | **Proliferating cells** | | |
| --- | --- | --- | --- | --- | --- | --- | --- | --- |
| **Category** | **No. of molecules** | **p-value range** | **Category** | **No. of molecules** | **p-value range** | **Category** | **No. of molecules** | **p-value range** |
| Cellular Development | 211 | 1.10E-05 - 3.18E-19 | Cell Death and Survival | 74 | 4.17E-03 - 7.83E-05 | Cellular Movement | 46 | 6.68E-03 - 2.64E-04 |
| Cellular Growth and Proliferation | 202 | 1.10E-05 - 3.18E-19 | Molecular Transport | 68 | 4.38E-03 - 3.40E-05 | Cellular Function and Maintenance | 44 | 6.68E-03 - 1.30E-06 |
| Cell Death and Survival | 177 | 1.32E-05 - 6.32E-12 | Small Molecule Biochemistry | 62 | 4.38E-03 - 2.22E-07 | Small Molecule Biochemistry | 36 | 6.68E-03 - 4.06E-05 |
| Cellular Movement | 158 | 1.43E-05 - 8.89E-17 | Carbohydrate Metabolism | 50 | 3.36E-03 - 2.22E-07 | Carbohydrate Metabolism | 19 | 6.33E-03 - 4.26E-09 |
| Lipid Metabolism | 99 | 1.64E-05 - 1.51E-11 | Lipid Metabolism | 34 | 4.38E-03 - 6.79E-06 | Cell Cycle | 7 | 6.68E-03 - 2.64E-04 |

**Suppl Table 3. Differentially expressed genes associated with uterine anomalies in the epithelial cell cluster. Excel file**

**Suppl Table 4. Differentially expressed genes associated with uterine anomalies in the stromal cell cluster. Excel file**

**Suppl Table 5. Differentially expressed genes associated with uterine anomalies in the proliferating cell cluster. Excel file**
